# Targeting Staufen 1 with antisense oligonucleotides for treating ALS and SCA2

**DOI:** 10.1101/2022.11.16.516816

**Authors:** Daniel R. Scoles, Sharan Paul, Warunee Dansithong, Karla P. Figueroa, Mandi Gandelman, Feliks Royzen, Collin J. Anderson, Stefan M. Pulst

## Abstract

Staufen1 (STAU1) is a multifunctional RNA binding protein that controls mRNA degradation and subcellular localization. STAU1 interacts with the ATXN2 protein, that is polyglutamine expanded in spinocerebellar ataxia type 2 (SCA2). We previously showed that STAU1 is elevated and aggregated in cells from SCA2 patients, cells from amyotrophic lateral sclerosis (ALS) patients, and in SCA2 and ALS mouse models. We also found that reduction of STAU1 abundance *in vivo* by genetic interaction improved motor behavior in an SCA2 mouse model, normalized the levels of several SCA2-related proteins, and reduced aggregation of polyglutamine-expanded ATXN2. Here we developed antisense oligonucleotides (ASOs) lowering STAU1 expression toward developing a therapeutic that may be effective for treating SCA2 and ALS. We performed a screen of 118 20mer phosphorothioate 2’-*O*-methoxyethyl (MOE) ASO gapmers targeting across the *STAU1* mRNA coding region for lowering STAU1 expression in HEK-293 cells. ASO hits lowering STAU1 by >45 % were rescreened in SCA2 patient fibroblasts, and 10 of these were tested for lowering STAU1 abundance *in vivo* in a new BAC-STAU1 mouse model. This identified efficacious ASOs targeting human *STAU1 in vivo* that normalized autophagy marker proteins, including ASO-45 that also targets mouse *Stau1*. When delivered by intracerebroventricular (ICV) injection, ASO-45 normalized autophagy markers and abnormal mRNA abundances in cerebella of ATXN2-Q127 SCA2 mice, as well as ChAT, NeuN and cleaved caspase-3 in spinal cord of *Thy1-* TDP-43 transgenic mice. Targeting *STAU1* may be an effective strategy for treating ALS and SCA2 as well as other disorders characterized by its overabundance.

## INTRODUCTION

STAU1 is an RNA binding protein that functions in RNA degradation and mRNA trafficking in neurons as well as nuclear exit of mRNA. We identified STAU1 as an interacting protein to ATXN2, a protein that is polyglutamine expanded in spinocerebellar ataxia type 2 (SCA2) (1). Because lowering *ATXN2* expression extends survival of ALS mice (2), we sought to investigate STAU1 in both SCA2 and ALS - relevant cells and mouse models to understand if STAU1 might be a viable therapeutic target for these disorders. We found that STAU1 is overabundant in SCA2 and other neurodegenerative diseases, including in the relevant cellular and transgenic mouse models and human patient tissues and cultured fibroblast cells. In addition to SCA2 these include Huntington’s disease (HD), C9orf72 ALS, and TDP-43 ALS (1,3,4).

STAU1 overabundance is associated with abnormal autophagy in multiple human tissues and mouse models of neurodegenerative disease (1,3,4). Mechanistically, this occurs by STAU1 direct interaction with the *MTOR* mRNA 5’-UTR resulting in its increased translation (4). Reducing STAU1 abundance normalized autophagy marker proteins in various systems including HEK-293 cells edited to express polyglutamine expanded ATXN2-Q58, and cultured patient fibroblasts from SCA2, TDP-43 and C9orf72 ALS patients, and transgenic mouse models with *TDP-43* or *C9orf72* mutations (3,4). Additionally, lowering Staufen abundance improves SCA2 mouse motor phenotypes and protein aggregations and mTOR-related autophagy phenotypes (1,3,4).

Our prior work on staufen suggests that lowering *STAU1* expression might be therapeutic for SCA2 and ALS. Here we present the development of an antisense oligonucleotide (ASO) therapeutic targeting *STAU1*. Among our findings are additional evidence that lowering *STAU1* modifies the mTOR-related autophagy phenotype in cultured cells and in a BAC-STAU1 transgenic mouse model that we developed. In this study we also identified ASOs targeting mouse *Stau1* useful for proof-of-concept studies in mice. We demonstrated improvement of neuronal phenotypes in SCA2 and TDP-43 transgenic mice treated with the *Stau1* ASO.

## MATERIALS AND METHODS

### Antisense oligonucleotides (ASOs)

ASOs used in this study were 20mer 5-10-5 2’-*O*-methoxyethyl (MOE) gapmers, phosphorothioated at all positions. All cytosine positions were methylated. ASOs were produced and purified by Integrated DNA Technologies (IDT). Preparations included desalting for *in vitro* work or HPLC purification and dialysis to remove chromatography solvent for *in vivo* work.

### Cell culture

HEK-293 or SCA2 patient-derived skin fibroblasts (SCA2-500-1), expressing ATXN2 q22/q45, were cultured and maintained in Dulbecco’s modified Eagle’s medium (DMEM, high glucose, ThermoFisher #11965118) supplemented with 10% fetal bovine serum, and 1x penicillin and streptomycin. The SCA2-500-1 patient fibroblasts were previously collected by us, with written consent, and the studies were approved by the Institutional Review Board at the University of Utah. HEK-293 cells were transfected with ASOs using Lipofectamine, while SCA2-500-1 cells were transfected with Lipofectamine 3000, according to manufacturer’s protocol (ThermoFisher). The cells were harvested at 48 or 72 hrs post-transfection for analyses.

### Cell Line Authentication

In order to adhere with the National Institutes of Health (NIH) guideline on scientific rigor in conducting biomedical research (NOT-OD-15-103) on the use of biological and/or chemical resources, we authenticated both the HEK-293 and SCA2-500-1 cell lines by STR analysis on 24 loci, including amelogenin for sex identification. The kit used for this was the GenePrint 24 system (Promega).

### Mice

ATXN2-Q127 (*Pcp2*-ATXN2[Q127]) (5) and B6;SJL-Tg(Thy1-TARDBP)4Singh/J (Jackson Laboratories; Stock 012836) (6) mouse lines were maintained as previously described (1). The BAC-STAU1 mouse model was developed by the University of Utah Transgenic Mouse Core by pronuclear microinjection of a non-linearized human Staufen1 bacterial artificial chromosome (BAC) construct (BAC-STAU1 clone RP11-120I11, BACPAC Resources) into fertilized oocytes sourced from mice with a B6/D2 mixed hybrid background (B6D2F1J, The Jackson Laboratory stock #100006). Prior to injection, the BAC construct was separated from other nucleotide fragments by pulsed field gel electrophoresis, and gel purified, by the University of Michigan Transgenic Mouse Core. Genotyping of mouse tail DNA was initially done with multiplex PCRs encompassing 17 PCR primer pairs covering all coding exons, 5’ and 3’ UTR and the 5’ upstream promoter region. Transgenic mouse bone marrow samples were sequenced and analyzed by Cergentis to determine integration sites and vector-vector breakpoints that represent concatemerization. BAC-STAU1 mice were maintained in a B6D2 mixed background by backcrossing to B6D2F1J no less than every 4 generations. The *Stau1^tm1Apa(-/-)^* (*Stau1^-/-^*) mouse (7) was a generous gift from Prof. Michael A. Kiebler, Ludwig Maximilian University of Munich, Germany, and maintained in a C57BL/6J background. Mice were maintained in a temperature and humidity-controlled environment on a 12h light/dark cycle with light onset at 6:00 AM. All studies and procedures were approved by an Institutional Animal Care and Use Committee (IACUC) at the University of Utah, SLC, UT.

### ICV ASO Injections

ASOs were delivered to mice by intracerebroventricular (ICV) injection using a Hamilton 26s gauge needle. Injections volumes were 10 μl of ASO diluted in phosphate buffered saline (PBS). Control mice received the same volume of normal saline. Injections were made under anesthesia with a mixture of oxygen and isoflurane, using a Stoelting stereotaxic frame. Anesthesia was initiated using 3% isoflurane for 5 min and the isoflurane mixture was lowered to 2% during injections. Stereotaxic bregma coordinates were −0.46 mm anteroposterior, −1.0 mm lateral (right side); −2.5 mm dorsoventral. Needles were removed 4 min after ASO delivery. Mice were maintained on a 39°C isothermal pad while anesthetized and during recovery.

### Quantitative PCR (qPCR)

For medium throughput assays performed in ASO screens and IC50 studies we used the Cells-to-Ct assays (ThermoFisher). These assays were combined with Taqman qPCR kits for human *STAU1* (ThermoFisher Hs00244999_m1) and human *ACTB* (Thermofisher Hs01060665_g1). For other qPCRs, standard SYBR Green assays were performed as follows: Total RNA was extracted from cultured cells, cerebellar tissues or spinal cord tissues using the RNeasyMini Kit (Qiagen Inc., USA) according to the manufacturer’s protocol. DNAse I treated RNAs were used to synthesize cDNAs using the ProtoScript cDNA First Strand cDNA Synthesis Kit (New England Biolabs Inc., USA). Primers for RT-PCR were designed to prevent amplification from genomic DNA (annealing sites in different exons or across intron-exon boundaries). Quantitative RT-PCR was performed in Bio-Rad CFX96 (Bio-Rad Inc., USA) with the Power SYBR Green PCRMasterMix (Applied Biosystems Inc, USA). PCR reaction mixtures contained SYBR Green PCRMasterMix and 0.5 pmol primers and PCR amplification was carried out for 45 cycles: denaturation at 95 °C for 10 sec, annealing at 60 °C for 10 sec and extension at 72 °C for 40 sec. The threshold cycle for each sample was chosen from the linear range and converted to a starting quantity by interpolation from a standard curve run on the same plate for each set of primers. Gene expression levels were normalized to the *Actin* mRNA levels. Four human transcripts, primers included STAU1-F 5’-TCCTTGGTTTCAAAGTCCCG-3’ and STAU1-R 5’-ATTTTCATCCCCAGAGCCAG-3’; ACTB-F: 5’-GAAAATCTGGCACCACACCT-3’ and ACTB-R: 5’-TAGCACAGCCTGGATAGCAA-3’. For mouse transcripts, primers included Stau1-F: 5’-AGTACATGCTCCTTACAGAACG-3’ and Stau1-R: 5’-TGATGCCCAACCTTTACCTG-3’; Aif1-A1: 5’-CTGGAGGGGATCAACAAGCAATTC-3’ and Aif-B2: 5’-CCAGCATTCGCTTCAAGGACATAA-3’; Gfap-for: 5’-CGGAGACGCATCACCTGTG-3’ and Gfap-rev: 5’-AGGGAGTGGAGGAGTCATTCG-3’; Actb-F: 5’-CGTCGACAACGGCTCCGGCATG-3’ and Actb-R: 5’-GGGCCTCGTCACCCACATAGGAG-3’; Mouse Actb-F: 5’-CGTCGACAACGGCTCCGGCATG-3’ and Actb-R: 5’-GGGCCTCGTCACCCACATAGGAG-3’.

### Western blot analyses

Protein extracts were prepared by homogenization of mouse cerebella in extraction buffer (25 mM Tris-HCl pH 7.6, 300 mM NaCl, 0.5% Nonidet P-40, 2 mM EDTA, 2 mM MgCl2, 0.5 M urea and protease inhibitors; Sigma; cat# P-8340) followed by centrifugation at 4°C for 20 min at 16,100 × g. Protein extracts were resolved by SDS-PAGE and transferred to Hybond P membranes (Amersham Bioscience Inc., USA). After blocking with 5% skim milk in 0.1% Tween 20/PBS, the membranes were incubated with primary antibodies in 5% skim milk in 0.1% Tween 20/PBS for 2 hrs at room temperature or overnight at 4°C. After washing in 0.1% Tween 20/PBS, the membranes were incubated with the corresponding secondary antibodies conjugated with HRP in 5% skim milk in 0.1% Tween 20/PBS for 2 hrs at room temperature and washed again. Signals were detected by using the Immobilon Western Chemiluminescent HRP Substrate (Millipore Inc., USA; cat# WBKLSO100) according to the manufacturer’s protocol, and detected using a ChemiDoc System (Bio-Rad). The intensity of proteins was determined using the ChemiDoc software and proteins were quantitated as a ratio to β-Actin.

### Antibodies

Antibodies included the following: rabbit polyclonal anti-Staufen antibody (Novus Biologicals, NBP1-33202); rabbit polyclonal anti-mTOR antibody (Cell Signaling Technology, 2972); rabbit polyclonal anti-Phospho-mTOR (Ser2448) [(1:3000), Cell Signaling Technology, 2971]; rabbit polyclonal anti-SQSTM1/p62 antibody (Cell Signaling Technology, 5114); rabbit polyclonal anti-LC3B antibody (Novus Biologicals, NB100-2220); mouse monoclonal anti-TDP-43 (human specific) antibody (Proteintech 60019-2-Ig); rabbit polyclonal human/mouse anti-TDP-43 antibody (Proteintech 10782-2-AP); mouse monoclonal anti-Calbindin-D-28K antibody [(1:5000), Sigma-Aldrich, C9848]; rabbit polyclonal anti-RGS8 antibody [(1:5000), Novus Biologicals, NBP2-20153]; mouse monoclonal anti-PCP2 antibody (F-3) [(1:3000), Santa Cruz, sc-137064]; rabbit polyclonal anti-PCP4 antibody [(1:5000), Abcam, ab197377]; Phospho-p70 S6 Kinase (Thr389) antibody [(1:3000), Cell Signaling, Cat# 9205]; GFAP (GA5) mouse mAb [(1:7000), Cell signaling, Cat #3670]; ChAT (E4F9G) Rabbit mAb [(1:5000), Cell Signaling, Cat# 27269]; NeuN (D4G4O) XP^®^ Rabbit mAb [(1:5000), Cell Signaling, Cat# 24307]; GAPDH (14C10) rabbit mAb [(1:7,000), Cell Signaling, Cat# 2118]; Cleaved Caspase-3 (Asp175) (5A1E) rabbit mAb [(1:3000), Cell Signaling, Cat #9664]; Anti-FAM107B antibody [(1:5,000) Abcam, ab175148); and mouse monoclonal anti-β-Actin-peroxidase antibody (clone AC-15) (Sigma-Aldrich, A3854). Secondary antibodies included peroxidase-conjugated horse anti-mouse IgG (Vector Laboratories, PI-2000) and peroxidase AffiniPure goat anti-rabbit IgG (Jackson ImmunoResearch Laboratories, 111-035-144).

### Immunohistochemical staining

Spinal cords were removed, and fixed in 4% paraformaldehyde/PBS and cryoprotected stepwise in 10, 20, then 30% sucrose/PBS. Tissues were embedded in OCT and sectioned (20 μm). Sections were labeled with human-specific anti-TDP-43 antibody or pan-TDP-43 antibody (see above) as previously described (1), and brightfield images were produced using an Olympus BX53 inverted microscope with a 40x objective. Regions of interest (ROIs) were selected manually around ventral horn MNs with clear evidence of a nucleolus. Staining intensity in ROIs was determined using NIH Elements software package.

### Behavioral phenotype testing

Mice were tested on the accelerating rotarod as previously described (8). Open field behavior was tested using a force plate actometer (9). Actometer hardware was as previously described (10,11), with center of mass tracked at 1000 Hz and saved using LabVIEW (National Instruments). We wrote custom code in MATLAB (MathWorks) to filter center of mass with a running average of 1 second, and then calculated the total distance traveled throughout the duration of the recording. The recordings were made in a small behavioral recording room, with animals placed in the room to habituate at least 30 minutes prior to recording. This room was well isolated from external noise, with dim, but equal lighting throughout the room as measured using a standard luxometer. Open field recordings were made over intervals of either 10 min, 20 s or 30 min, 20 s, allowing 20 s for the experimenter to leave the room and close the door after placing the animal in the chamber, and the total distance traveled was calculated for the specified duration beginning after 20 s.

### Electrophysiology

The preparation of parasagittal cerebellar slices closely followed our previously published description of the process (8). Cerebella from 10-week-old ATXN2-Q127 and WT littermates were removed and quickly immersed in 4 °C sucrose solution bubbled with 95% O_2_ and 5% CO_2_ (225 mM sucrose, 25 mM NaHCO_3_, 20 mM D-glucose, 3.0 mM MgCl_2_, 2.5 mM KCl, 1.25 mM NaH_2_PO_4_ and 0.5 mM CaCl_2_). Parasagittal cerebellar slices (400 μm) were sectioned using a vibratome (Vibratome 1000). Extracellular recordings were acquired in voltage-clamp mode at physiological temperature (34 ± 1°C) using a dual channel heater controller (Model TC-344B, Warner Instruments) and constantly perfused with carbogen-bubbled extracellular solution (119 mM NaCl, 26 mM NaHCO_3_, 11 mM D-glucose, 2.5 mM KCl, 2.0 mM CaCl_2_, 1.5 mM MgCl_2_ and 1.0 mM NaH_2_PO_4_) at a rate of 3 mL per minute. Cells were visualized on an upright microscope (Zeiss Axioskop 2) with a 40x water-immersion lens. Borosilicate glass pipettes with resistances between with 1 to 3 MΩ were filled with extracellular solution and used for recording action potential-associated capacitive current transients. The pipette potential was held at 0 mV and placed close to the Purkinje neuron axon hillock (soma/axon). Data were acquired at 20 kHz using a Multiclamp 700B amplifier, Digidata 1440 with pClamp10 (Molecular Devices) and bandpass filtered between 0.1 to 5 kHz. Each Purkinje neuron recording spanned a duration of 2 minutes and an average of 43 cells were measured from each mouse. Each treatment group contained approximately 5 mice that were injected at 8 weeks of age with ASO or PBS. Data were analyzed with custom MATLAB scripts (MathWorks) and Prism (GraphPad).

### Statistical Analysis

Statistical differences between selected groups evaluated by qPCR, western blotting, and the electrophysiological data were also evaluated using analysis of variance (ANOVA) tests followed by post-hoc tests of significance (Tukey’s or Bonferroni’s tests for multiple comparison, as indicated). Grip strength data were tested using repeated measures ANOVA. Student’s *t*-tests were unpaired and two-tailed. All tests were performed using the GraphPad Prism software package except for tests of open field behavior data that were performed in MATLAB.

## RESULTS

### Identification of lead *STAU1* ASOs

Our ASO screen began with an *in silico* design stage where we determined all possible 20 base pair “k-mers” from the start of the 5’-UTR through the end of the 3’-UTR, for a total of 3664 ASO sequences. ASOs with sequences predicted to reduce efficacy were eliminated. ASOs with CpGs were eliminated to reduce immunoreactivity. Any remaining cytosines were 5’-methylated. ASOs with 3 or more consecutive Gs were excluded to avoid G quadraduplexing. Only sequences with ΔGs > −10 for hairpinning were allowed. The %GC content was limited to 40-60% for optimal annealing. We also eliminated ASOs not aligning with cynomolgus sequences for later NHP work. Of all ASOs meeting these criteria, we selected 118 ASOs spaced ~ 10 bp apart, and synthesized them as fully phosphorothioated (PS) 5-10-5 2’-*O*-methoxyethyl (MOE) gapmers. To determine the efficacy for an ASO with sequence identity to mouse and human Staufen that may be useful for *in vivo* work, we included ASO-45 in our screen despite its sequence did not strictly conform to our conservative criterion in that it has a tract of 3 consecutive Gs and a CpG dinucleotide. Of all the ASOs in our screen, only one other also had sequence identity to mouse *Stau1*, ASO-308. We then screened the 118 ASOs in HEK-293 cells transfected at a single 50 nM dose in duplicate, and performed qPCR to determine *STAU1* mRNA abundance following 48 hours transfection time. ASO-249 was the most efficacious ASO, with 38% STAU1 mRNA abundance remaining, and ASO-045 was the third-most efficacious ASO (Fig. 1A).

**Fig. 1.**
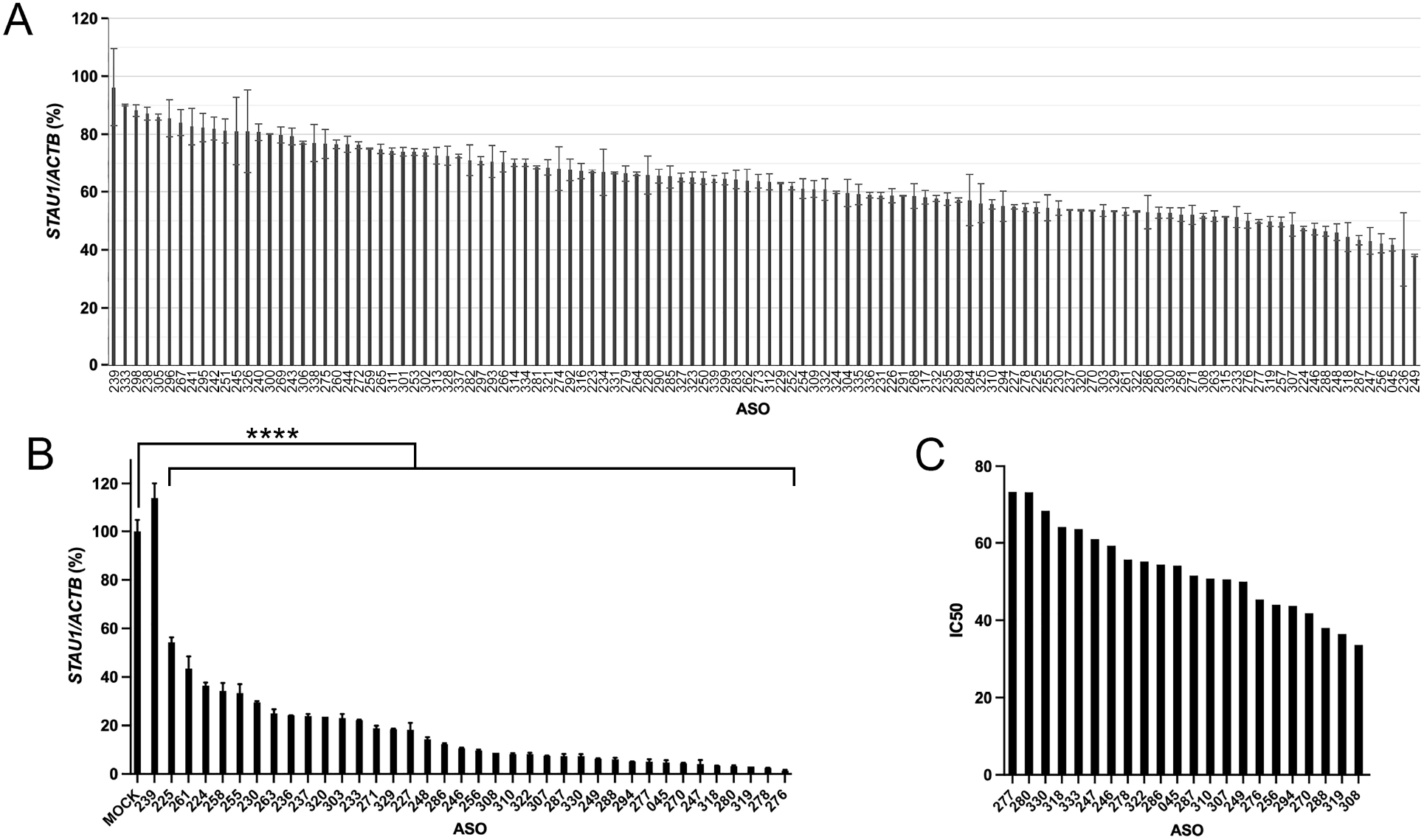
ASO screen. A) 118 STAU1 ASOs from the *in silico* design step were screened in HEK-293 cells in duplicate transfections at 50 nM. The *STAU1/ACTB* values shown are relative to mock transfected HEK-293 set to 100%, and are mean ± SD for duplicate wells. B) A selection of efficacious *STAU1* ASOs from the primary screen were rescreened in SCA2 (Q22/Q45) patient fibroblasts in triplicate at 100 nM. The least effective ASO in (A), ASO-239, was included as a negative control, and did not lower *STAU1* expression in SCA2 FB cells. Values are mean ± SD. All pairwise comparisons to the mock transfected control are significantly different (****, p<0.0001), determined by one-way ANOVA with the Tukey correction for multiple comparisons. C) IC50s from 6-point concentration curves compared for the 22 ASOs. Charts from which IC50s were derived are shown in Supplemental Data Fig. 1. In all cases, cellular *STAU1* mRNA levels were determined by qPCR after 48 hours transfection time.

The next task was to screen a selection of the most efficacious ASOs from the HEK-293 screen in SCA2 (Q22/Q45) patient fibroblasts. The most efficacious 37 ASOS were screened in the SCA2 patient fibroblasts, along with ASO-239 that was least effective in HEK-293 cells as a negative control. Cells were transfected at 100 nM in triplicate for 48 hours. The maximum *STAU1* reduction was 98.6% for ASO-276 and ASO-045 ranked 8th (Fig. 1B).

We then determined dose response curves for the most efficacious 22 ASOs for lowering *STAU1* mRNA expression in SCA2 patient fibroblasts. Assays were performed in batches of ASOs, including ASO-045 in each batch as a comparative control to assess reproducibility. Six ASO doses were included with each dose triplicated. We observed remarkable reproducibility among five batch sets, with the IC50 standard deviation for ASO-045 varying by only ± 2.2 nM (Supplementary Data Fig. 1). Comparison of the IC50 values among the 22 most efficacious ASOs revealed ASO-308 as the most potent of all the ASOs (Fig. 1C).

We then aligned each of the most efficacious ASOs with the human GRCh38/hg38 genome identifying potential off targets for some ASOs. ASO-308 and ASO-287 have one and two mismatches to *SORL1*, respectively, that has loss of function mutations in Alzheimer’s disease (12). ASO-294 has 1 mismatch to *FUT8* and *PARD3B* that are both expressed in astrocytes by querying Brain RNA-seq (13). ASO-278 that has one mismatch to *POLG2* that is mutated in epilepsy (14). These ASOs were not considered further with the exception of ASO-308 use in wildtype mice, below. We also aligned ASO-45 to the mouse MM10 genome revealing no potential off targets in the mouse.

To begin to characterize efficacy for the 10 most potent ASOs for normalization of pathologically elevated STAU1 we utilized CRISPR-Cas9 edited HEK-293 cells expressing polyglutamine expanded ATXN2 (ATXN2-Q58 KI cells), that we have previously demonstrated have elevated STAU1 abundance (1,3). We confirmed significantly elevated STAU1 in ATXN2-Q58 KI cells compared to HEK-293 cells with unmodified ATXN2 and transfection of each of these 10 ASOs significantly lowered STAU1 levels to quantities equal to that in unmodified HEK-293 cells (Fig. 2).

**Fig. 2.**
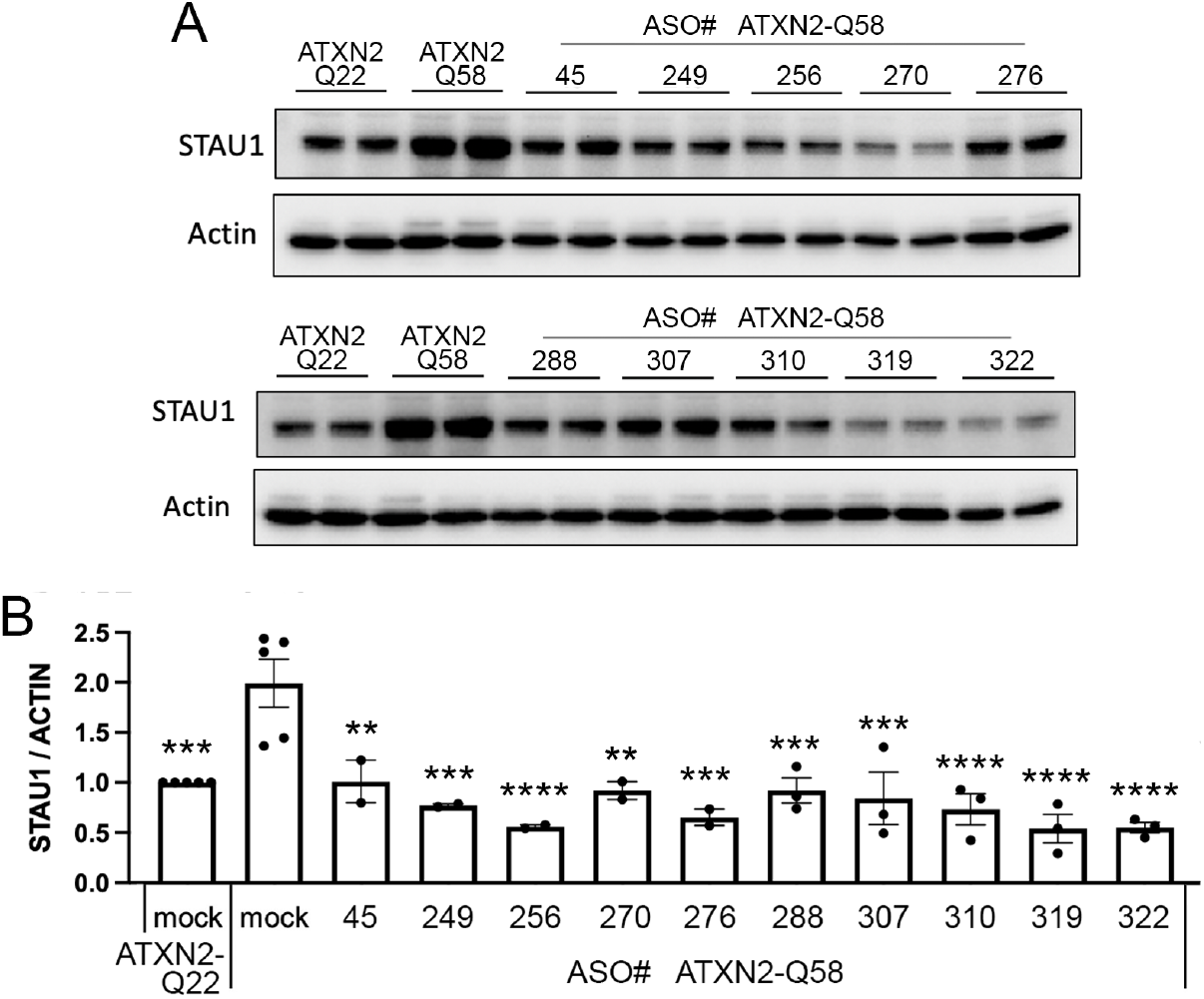
Effects of ASOs for lowering pathologically elevated STAU1 abundance and on normalizing autophagy-related proteins. A) HEK-293 ATXN2-Q58 KI cells that have elevated STAU1 abundance relative to non-mutant HEK-293 ATXN2-Q22 cells were transfected in duplicate cultures with 100 nM of each of 10 lead ASOs. Following 48 hrs, STAU1 levels was determined in cell lysates by western blotting. B) Quantifications of the blots shown in (A). Values are means ± SD relative to Actin. Blots were performed in duplicate or triplicate.

### Generation of BAC-STAU1 mice and *in vivo* ASO screening

Evaluation of ASOs *in vivo* was made possible by the production of a BAC mouse harboring the human *STAU1* gene. We utilized a 133.8 kDa BAC construct (Fig. 3A) and produced transgenic mice by pronuclear injection. Sequencing of BAC-SCA2 mouse bone marrow specimens determined the integration sites of 8 potential founder lines. Of these, only one founder, BAC-STAU1.6, had a single genomic integration of the transgene, on chromosome 17 (chr17:9,394,829-~9,610,924). The integration occurred simultaneously to a 210 kb genomic deletion within this region. By querying the NCBI Reference Sequence Database (RefSeq), there are no annotated genes in this region of the mouse genome and therefore no gene disruption is predicted. No structural variants were identified within the integrated construct. Due to the nature of the sequence at the integration site, the copy number via this method could not be estimated.

**Fig. 3.**
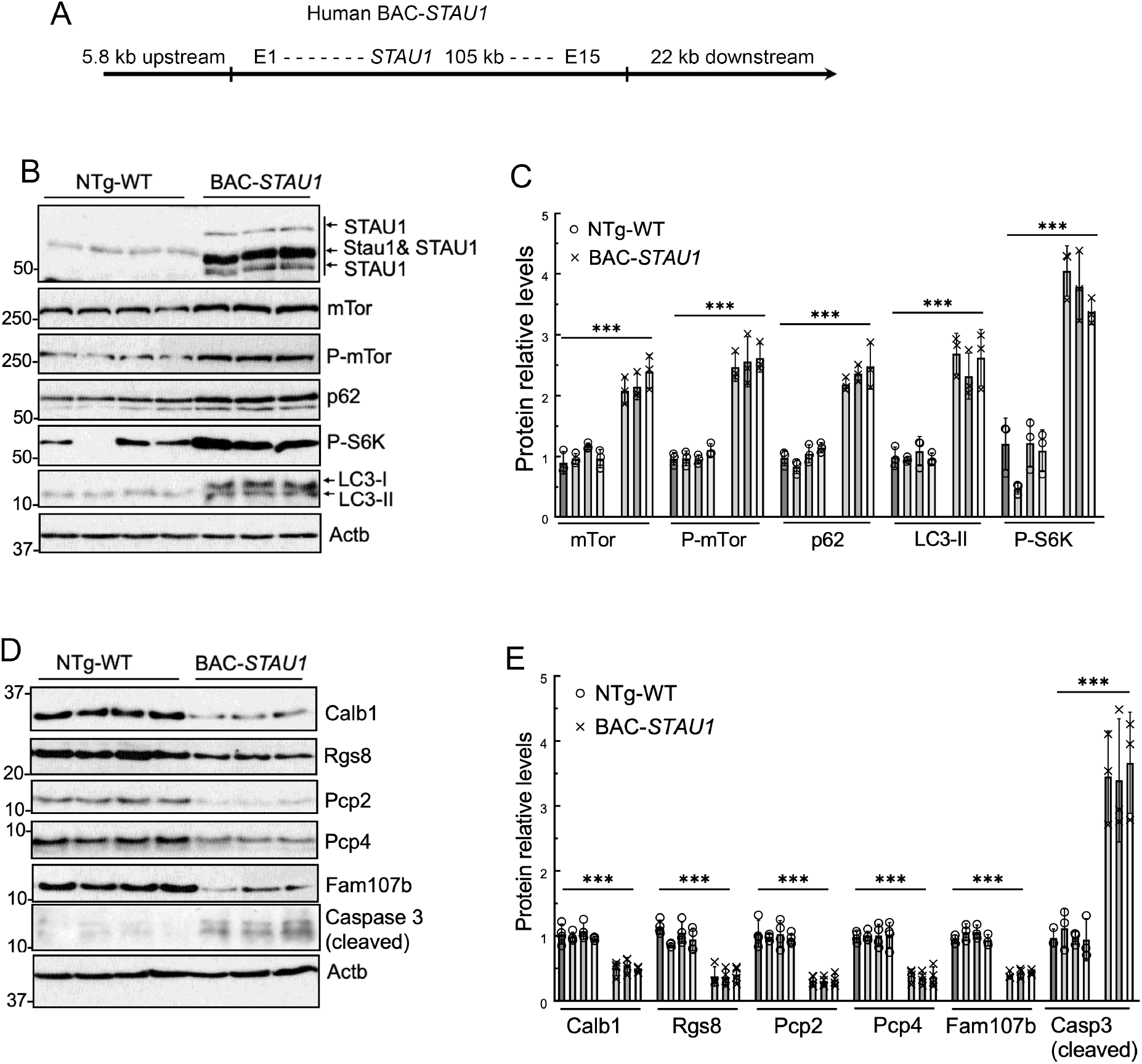
Generation and characterization of a *BAC-STAU1* transgenic mouse model. A) Schematic of the BAC construct showing the relative position of the *STAU1* gene and flanking sequences. B-E) Western blotting showing autophagy and other abnormal molecular phenotypes in the cerebellum of the *BAC-STAU1* mouse model. B) BAC-*STAU1* mouse cerebellar extracts (14 weeks of age; 3-4 animals per group) showing expression of STAU1 (3 isoforms), abundance of mTOR, phospho-mTOR, phospho-S6K, p62 and LC3-II. C) Quantification of B. D) *BAC-STAU1* mice had decreased levels of CALB1, RGS8, PCP2, PCP4 and FAM107b, and increased cleaved caspase 3 in cerebellar extracts. E) Quantification of D). Each lane represents an individual mouse. β-Actin was used as a loading control.

BAC-STAU1 mice express an increased abundance of STAU1 protein in cerebellum that is associated with elevated and hyperphosphorylated mTOR as well as increased levels of other proteins in the autophagy signaling pathway including phospho-S6K, p62 and LC3-II (Fig. 3B,C). BAC-STAU1 mice also displayed reduced levels of *Calb1, Rgs8, Pcp2, Pcp4* and *Fam107b* in cerebellum (Fig. 3D,E).

We next evaluated efficacy for the 10 most potent ASOs in BAC-STAU1. We treated 8 wk old BAC-*STAU1* mice by intracerebroventricular (ICV) injection of 300 μg ASO for 2 wks, and evaluated cerebellar and spinal cord proteins on western blots. Each of the ASOs significantly reduced STAU1 abundance both on western blots (Fig. 4A–D). Additionally, for all ASOs the abnormally elevated levels of CALB1, GFAP, mTOR, p62, and LC3-II were markedly reduced or normalized in cerebellum (Fig. 4A–D). We also evaluated *STAU1, Aif1*, and *Gfap* in cerebellum and spinal cord by qPCR (Fig. 4E,F). Interestingly, some ASOs did not significantly reduce *STAU1* mRNA levels in cerebellum despite that they lowered STAU1 protein on western blots suggesting *STAU1* ASOs may be effective for inhibiting translation. Notable ASOs include ASO-256 and ASO-319 that were consistently potent against both STAU1 protein and *STAU1* mRNA and did not significantly elevate *Aif1* or *Gfap* abundance.

**Fig. 4.**
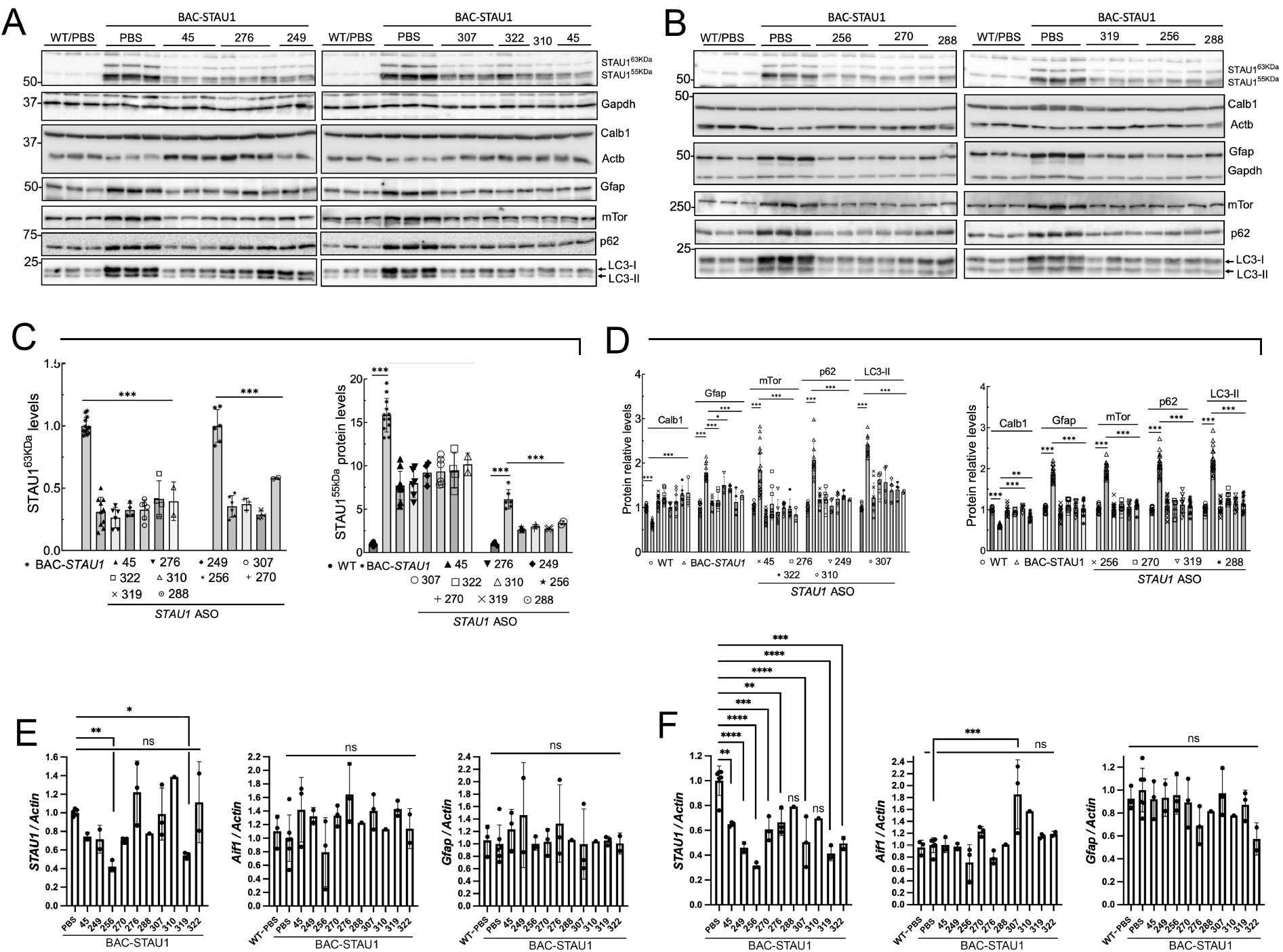
*In vivo* testing of the top 10 lead ASOs in BAC-SCA2 mice. BAC-STAU1 mice were injected ICV at 8 weeks of age with 300 μg ASO and evaluated by western blotting and qPCR following 2 weeks treatment time. A-D) Western blotting. Shown are quantified western blot data for STAU1 and other proteins that we previously shown modified with STAU1 expression (CALB1, GFAP, mTOR, p62, LC3-II) that are improved or restored following treatment with the ASOs. A) ICV injection set 1 including ASOs 045, 276, 249, 307, 322, and 310. B) ICV injection set 2 including ASOs 256, 270, 319, and 288. C) Quantification of the 63 kDa and 55 kDa forms of STAU1 on blots shown in A & B. D) Quantification of CALB1, GFAP, mTOR, p62, LC3-II on blots shown in A & B. Values are means ± SD relative to Actin (for STAU1, CALB1, p62, LC3) or GAPDH (GFAP, mTOR). Blots were performed in duplicate or triplicate. E-F) qPCR: Shown are the mean ± SD *STAU1, Aif1*, and *Gfap* expression values relative to Actin for the indicated ASOs in cerebellum (E) and spinal cord (F). Values are mean ± SD. One-way ANOVA with Bonferroni corrected probabilities: *, p<0.05; **, p<0.01; ***, p<0.001; ****, p<0.0001; ns, not significant.

We then compared the efficacious ASOs in doses in *BAC-STAU1* mice. We selected ASO-256 and ASO-319 that were the two most efficacious ASOs for lowering Staufen1 protein and mRNA *in vivo* (Fig. 4A), and we included ASO-249 and ASO-270 as the next two most efficacious ASOs based on the *in vivo* data, also supported by efficacy in ATXN2-Q58 KI cells (Figs. 2 and 4). These four ASOs are among the top 8 ranked by IC50 (Fig. 1C). We further included ASO-45 that also targets mouse *Stau1*. In this experiment we treated 8 wk old *BAC-STAU1* mice with 300, 600 and 900 μg ASO ICV for 2 weeks. Acute toxicity was evident as the number of surviving animals was reduced with dose, with all animals lost that were injected with 900 μg ASO-249 or ASO-319.

We also assessed *Aif1* and *Gfap* for microgliosis and astrogliosis, respectively, and observed no significant increases except for ASO-256 that elevated the expression of both genes at higher doses (Fig. 4B,C). ASO-256 was eliminated form further consideration.

### ASOs targeting mouse *Stau1*

Our screen had produced two ASOs that target human *STAU1* that also have sequence identity to mouse *Stau1*, ASO-45 and ASO-308. To compare their efficacy, we treated 11 wk old wildtype mice with 500 μg ASO by ICV injection with 2 wk treatment time. At the endpoint we evaluated cerebellar proteins on western blots demonstrating ~70% reduction of Stau1 protein levels compared to PBS treated control mice (Fig. 5A,B). We also evaluated *Stau1, Aif1* and *Gfap* mRNA levels in cerebellum by qPCR. *Stau1* mRNA levels were ~40% reduced and no activation of *Aif1* was observed for either ASO. However, a significant 15% increase of *Gfap* was observed for ASO-308 (p=0.005). Based on this we performed *in vivo* proof-of-concept studies using ASO-45.

**Fig. 5.**
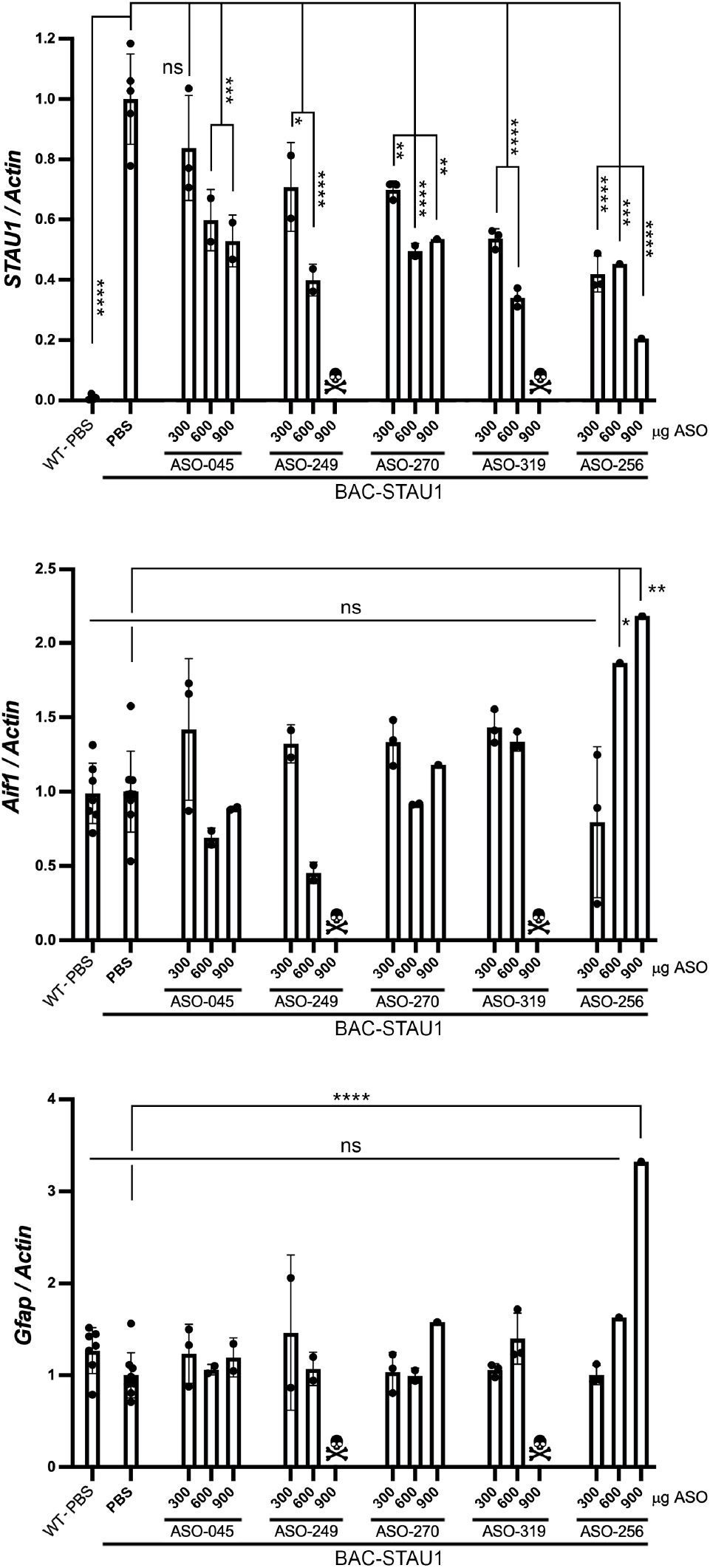
Dose testing for lead ASOs in BAC-STAU1 mice. BAC-STAU1 mice aged 8 weeks were treated with the indicated ASOs and doses. Following 2 weeks treatment cerebellar *STAU1, Aif1* and *Gfap* were determined by quantitative PCR. N=3 mice per dose. Points represent mean ± SD for the qPCR values per mouse. One-way ANOVA with Bonferroni corrected probabilities: *, p<0.05; **, p<0.01; ***, p<0.001; ****, p<0.0001; ns, not significant.

### Modification of SCA2 cerebellar Purkinje cell firing by ASO-45

Previously, we showed that cerebellar Purkinje cells in ATXN2-Q127 transgenic mice have reduced firing frequency that could be improved by lowering the expression of *ATXN2* with an ASO (8). We performed a similar experiment by treating 8 wk old BAC-STAU1 mice or wildtype littermate control mice with 500 μg ASO-45 or PBS vehicle by ICV injection. After two weeks treatment time we made extracellular recordings from PCs in acute cerebellar slices. As expected, we observed significantly reduced PC firing frequency in ATXN2-Q127 mice compared to the WT littermates and the PC firing frequency in ATXN2-Q127 mice was significantly improved following treatment with ASO-45 (Fig. 7A). The PC firing frequency in wildtype littermates was unaffected by the ASO treatment (Fig. 7A). Evaluation of target engagement showed that *Stau1* mRNA was significantly reduced in cerebellum of ATXN2-Q127 mice at the endpoint (Fig. 7B).

**Fig. 6.**
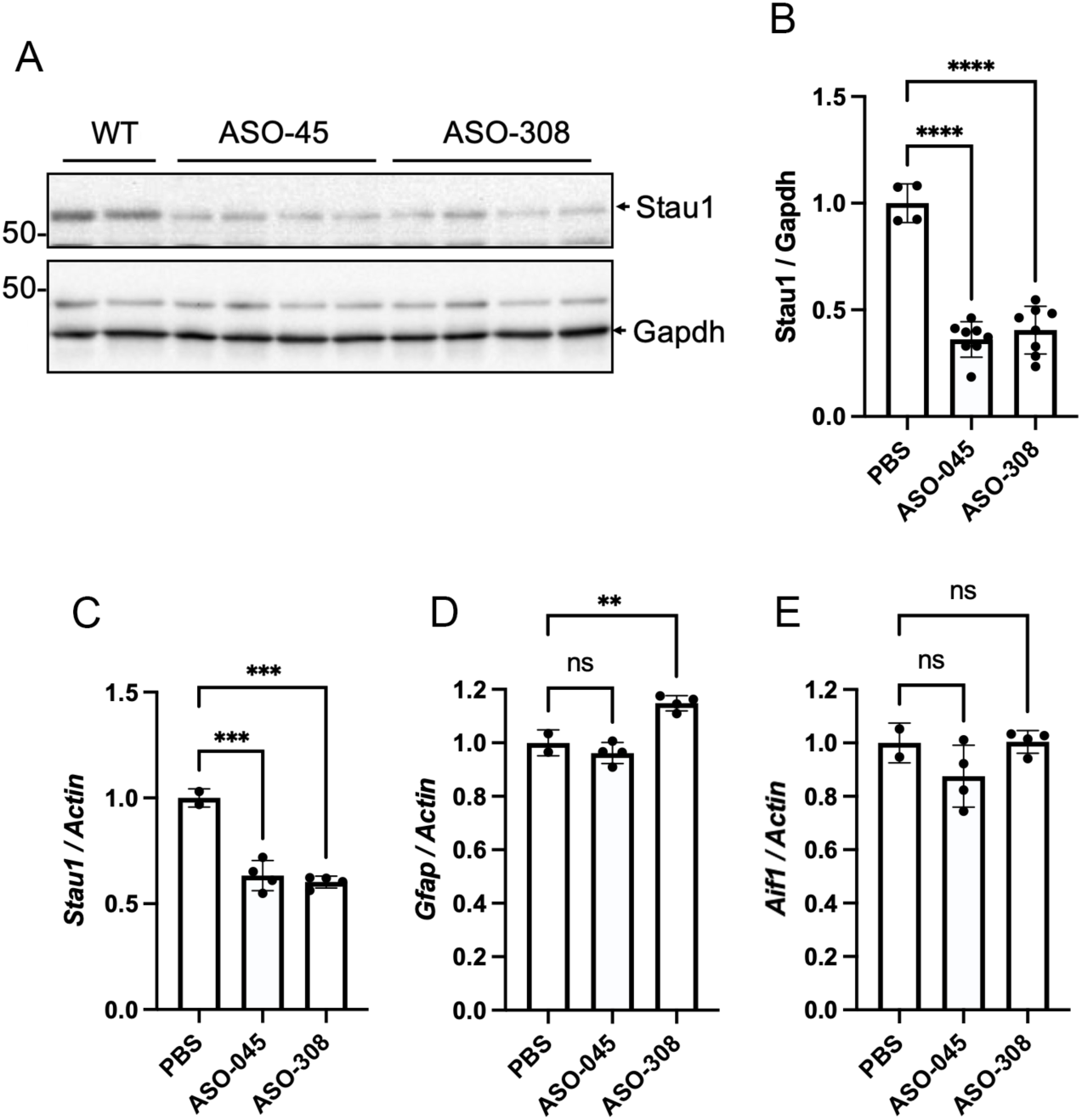
ASO-45 and ASO-308 target *Stau1* in wildtype mice. Wildtype mice were treated at 11 wks of age with 500 μg ASO, and were evaluated for molecular changes at 13 wks of age. N=2 WT injected with PBS and 4 WT mice injected with either ASO-045 or ASO-308. A&B) Western blotting of cerebellar protein extracts (A) and quantification (B). Each lane represents a different mouse, and the blot was replicated once. C-E) Determination of *Stau1* mRNA levels in cerebellar extracts by qPCR (C) with evaluation of *Gfap* (D) and *Aif1* (E), markers of cytotoxicity. One-way ANOVA with Bonferroni corrected probabilities: **, p<0.01; ***, p<0.001; ****, p<0.0001; ns, not significant.

**Fig. 7.**
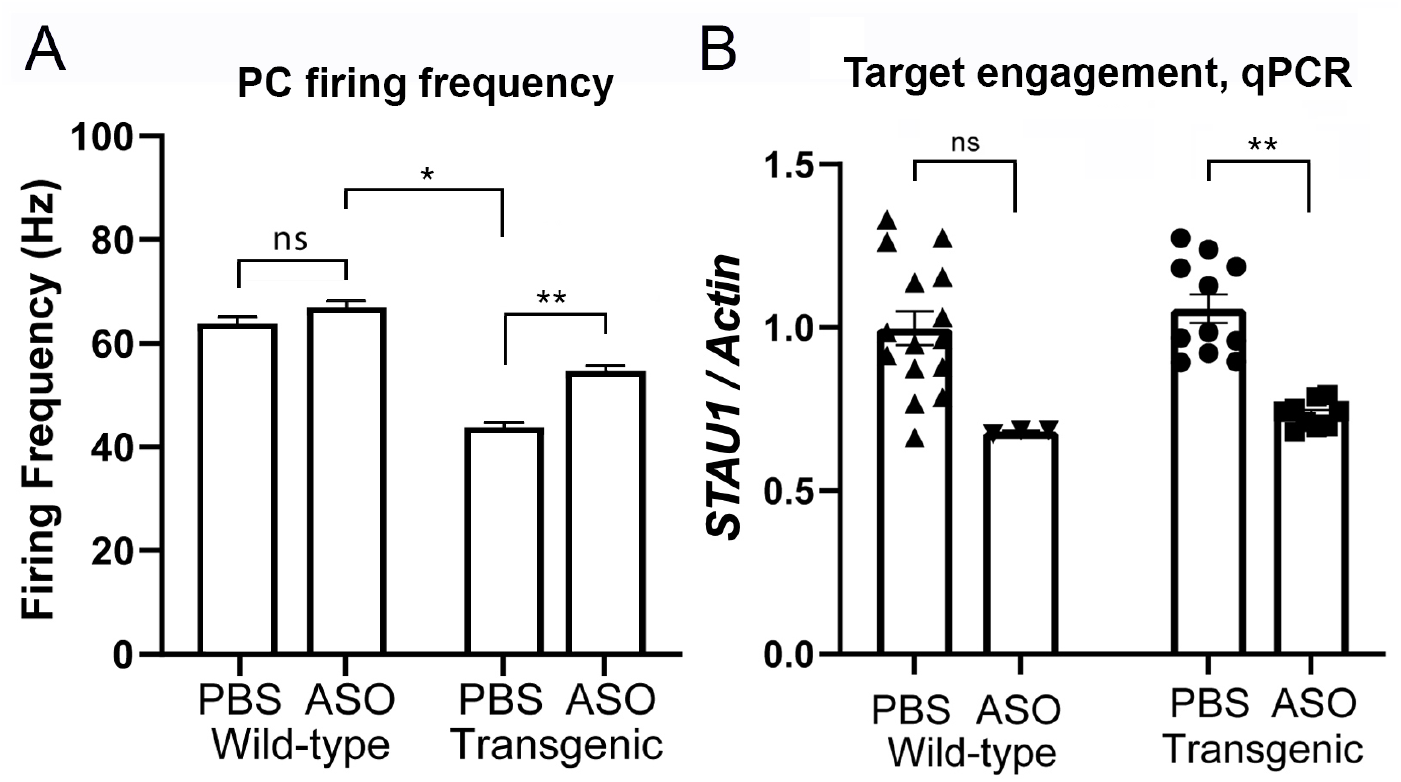
*Stau1* ASO-45 modifies Purkinje cell (PC) firing frequency in SCA2 ATXN2[Q127] mice. ATXN2[Q127] mice aged 8 weeks were treated with 500 μg of ASO-45 that targets mouse *Stau1* or PBS vehicle. A) Following 2 weeks treatment extracellular recordings were made from PCs in acute cerebellar slices. N=4-5 mice, 30-50 neurons recorded per mouse. Values are means ± SEM. B) Target engagement determined by qPCR. Statistical tests were performed by ANOVA using the mixed-effects model. *, p<0.05; **, p<0.01; ns, not significant.

### Normalization of spinal cord proteins in TDP-43 mice treated with ASO-45

We sought to determine abnormally abundant ChAT, NeuN and Gfap in older *Thy1*-TDP-43^Tg/+^ mice could be modified by treatment with ASO-45. We treated 24 wk old *Thy1*-TDP-43^Tg/+^ mice with ASO-43 by ICV injection for a duration of 2 weeks. At the endpoint we evaluated Stau1, human TDP-43, ChAT, NeuN and GFAP on western blots. We observed significant reductions of ChAT and NeuN that were each fully normalized in ASO-45 treated *Thy1*-TDP-43^Tg/+^ mice, associated with significant reduction of TDP-43 protein abundance (Fig. 8A,B). GFAP was also highly elevated in *Thy1*-TDP-43^Tg/+^ mice and was significantly reduced by ASO-45 treatment but the level of GFAP in treated mice did not return fully to normal (Fig. 8A,B). We also observed reduction of TDP-43 and ChAT in spinal cord motor neurons by quantitative immunohistochemical staining, that were improved after the ASO-45 treatment (Fig. 8B,C).

**Fig. 8.**
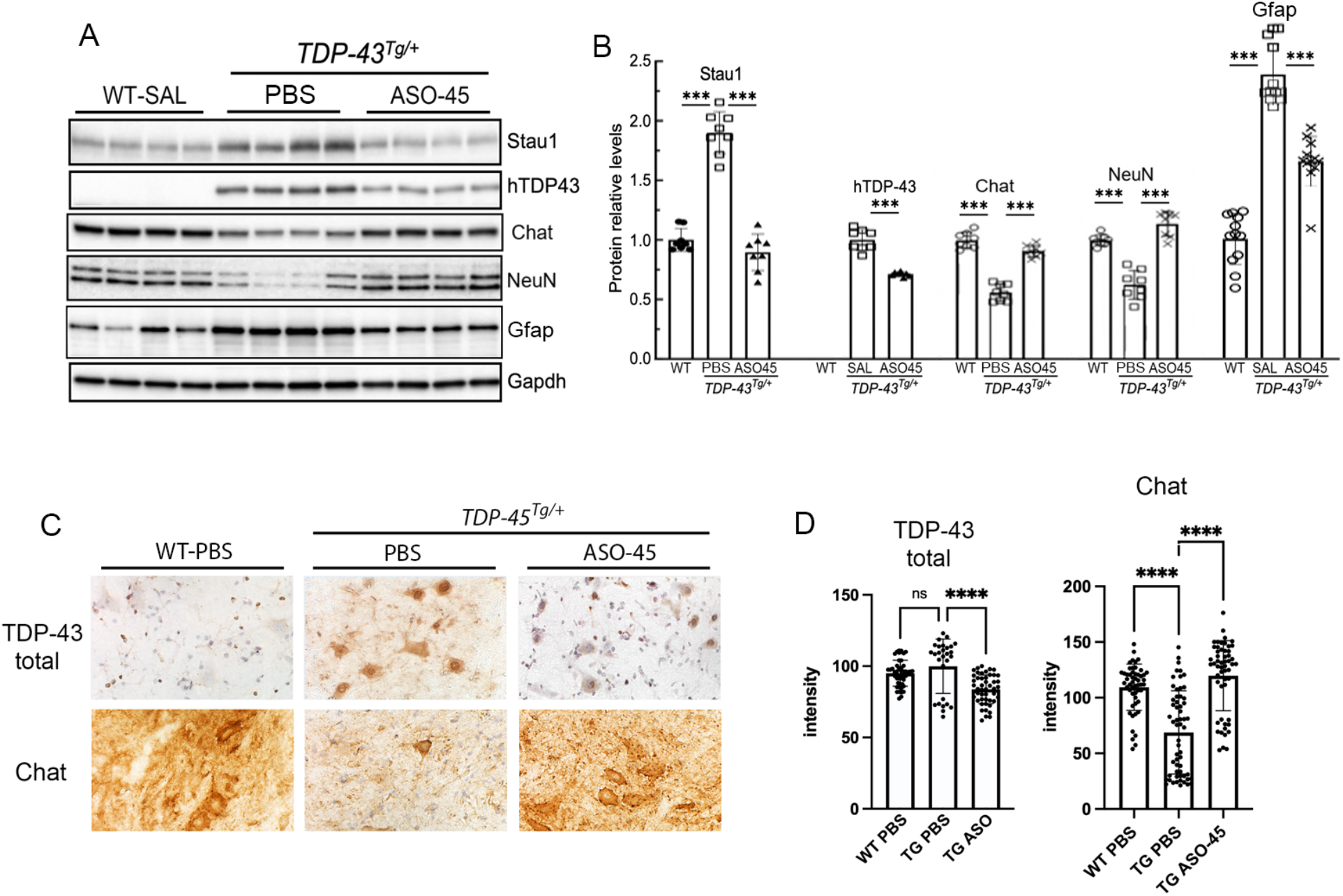
STAU1 ASO-45 normalized neuronal health markers in spinal cord of *Thy1*-TDP-43^TG/+^ transgenic mice. Mice 24 wks of age were treated with ASO-45 or PBS for 2 wks and proteins from total spinal cord extracts were evaluated by western blotting. Proteins determined were human TDP-43 (hTDP-43), ChAT, NeuN and GFAP relative to GAPDH (A). Quantification of two blots (B). C & D) Immunohistochemical staining for TDP-43 in spinal cord. C) TDP-43 staining increased in motor neurons of TDP-43^TG/+^ mice vs WT mice, and was reduced in TDP-43^TG/+^ mice treated with ASO-45. D) Quantification of TDP-43^TG/+^ staining intensity in MNs from sections across the spinal cord including cervical, thoracic and lumbar regions. Probabilities were determined by one-way ANOVA and were adjusted using the Bonferroni correction: *, p<0.05; ***, p<0.001; ****, p<0.0001.

### Investigating the on-target effects of lowering STAU1

Alterations of motor phenotypes were not previously observed in mice null of *Stau1*, except for mildly reduced open field locomotor activity (7). To confirm the observation, we performed open field behavioral testing using 6 month old *Stau1^+/-^*, *Stau1^-/-^*, and wildtype littermate mice in a force plate chamber. No significant reductions on open field behavior determined as distance travelled was observed (Fig. 9A). We also showed no significant difference in percentage of time in chamber center 50%. We also observed no sex differences. Recordings of *Stau1* knockout mice were made over 30 min. Analysis of the data indicated that sufficient power of the test would be obtained in 10 min recordings, that we employed for the study of ASO-45 on open field behavior. To determine how ASO-45 modified open field behavior we treated WT mice that were 3 months of age with 300 μg ASO-45 for a duration of 3 wks. Again, we observed no significant reduction in open field behavior, including distance travelled or time in chamber center 50%, and we observed no sex differences. At the endpoint we evaluated *Stau1* mRNA levels in cerebellum by qPCR demonstrating significant *Stau1* reduction in ASO-45 treated mice (Fig. 9C) to levels that were similar to reductions seen in Figs. 6 and 7 for ASO-45. We also performed a Pearson’s correlation analysis of distance traveled to *Stau1* mRNA abundance, demonstrating no significant correlation (Fig. 9D).

**Fig. 9.**
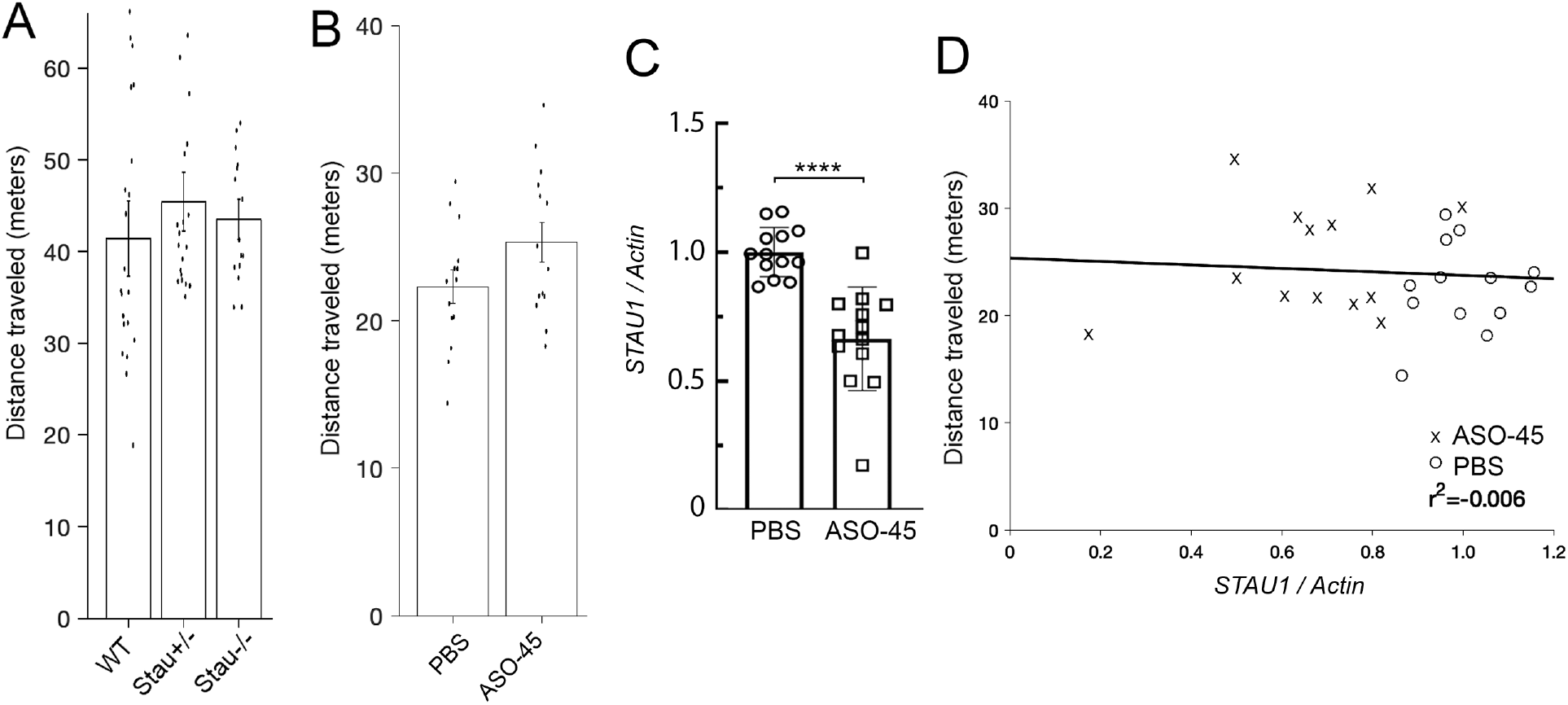
On target open field behavior testing for lowering *Stau1* in adult mice. A) Distance travelled for center of mass for *Stau1* heterozygous knockout (*Stau1^+/-^*), *Stau1* homozygous knockout (*Stau1^-/-^*) and wildtype littermate mice 6 months of age, recorded over 30 min. No significant difference was observed between groups for genotype or sex (ANOVA, p>0.05). N= 21 WT (10 males 11 females), 20 Stau1^+/-^ (10 males 10 females), 15 Stau1^-/-^ (8 males 7 females). B-D) WT mice age 3 months treated ICV with 300 μg ASO-45 for 3 wks. B) Distance travelled for center of mass recorded over 10 min (Student’s *t*-test, p=0.098). No significant difference was observed between sexes (p>0.05). N=14 mice (7 males 7 females) per group. C) Target engagement in brain by qPCR (Student’s *t*-test p<0.0001). D) Plot of *Stau1* mRNA abundance vs distance revealing no correlation. Pearson’s correlation test p=0.73, r=0.078, r^2^=-0.006. Values are means and SEM (A-B) or SD (C). Experiments in A & B showed no significant difference in percentage of time in chamber center 50%.

## DISCUSSION

### Development of disease modifying therapeutics

There is a critical need to develop disease modifying therapeutics for neurodegenerative disorders. At the present time there are only a handful of such therapeutics that are approved by the FDA for use in the clinic for movement disorders including spinal muscular atrophy (SMA) (nusinersen, onasemnogene abeparvovec and risdiplam) and Duchenne’s muscular dystrophy (DMD) (eteplirsen, golodirsen and viltolarsen) (15). Addressing this need are a number of ongoing preclinical studies and clinical trials on ASO therapeutics for neurodegenerative diseases (16). Among these therapeutics is BIIB105, an ASO that we developed that targets *ATXN2* (8) that is currently in Phase I clinical trial (NCT04494256) for ALS with or without CAG repeat expansions in *ATXN2*. The preclinical version of BIIB105, differing by its sequence, was effective for improving or normalizing cerebellar molecular, motor, and electrophysiological phenotypes of ATXN2-Q127 mice (8). A simultaneous study in which one of us (SMP) participated found the reduction of mouse *Atxn2* expression effective for improving motor, stress granule and survival phenotypes in *Thy1-TDP-43* ALS/FTD mice (2). We subsequently found that ATXN2 interacts with STAU1, an RNA-binding protein that we discovered was overabundant in CNS tissues of ATXN2-Q127 and *Thy1*-TDP-43 mice as well as skin cell fibroblasts from SCA2, HD, AD, and ALS patients (1,3). The overabundance of STAU1 and its established role in regulating mRNAs in stress granules pointed to it as a logical therapeutic target for neurodegenerative disease. Indeed, *STAU1* might be a preferred therapeutic target to *ATXN2* because of its overabundance, in particular when *ATXN2* is non-mutant.

### Studies supporting STAU1 therapeutics

Support for targeting *STAU1* in neurodegenerative disease came not only from our discovery of its overabundance in multiple patient cells and animal models, but also the associated abnormalities in the autophagy pathway that could be improved by lowering *STAU1* expression. Each of the patient fibroblasts in which we charactered overabundant STAU1 had hyperactivated mTOR and overabundance of autophagy marker proteins p62 and LC3-II. The same was true in the cerebella and spinal cords of ATXN2-Q127 mice and *Thy1*-TDP-43 mice (3). We were able to pinpoint the reason for these abnormalities, by demonstrating a direct interaction between STAU1 and the 5’-UTR of the *MTOR* mRNA transcript resulting in its increased translation (4). Likewise, we observed reduced expression of the cerebellar gene *PCP2* in SCA2 patient skin cell fibroblasts, previously confirmed in cerebella of ATXN2-Q127 mice (5,8), that we accounted for by STAU1 direct binding of the 3’-UTR of the *PCP2* mRNA transcript entering it into the Staufen1 mediated mRNA decay (SMD) pathway (1). These abnormal pathways of autophagy and PCP2 protein expression could be normalized by lowering *STAU1* expression by RNAi (1,3). STAU1 also bidirectionally modulates activation of the unfolded protein response (UPR). Overexpression of STAU1 caused pro-apoptotic activation of the UPR, evidenced as an increase in PERK, p-eIF2*α* and CHOP proteins. Conversely, in TDP-43 and C9ORF72 mutant patient-derived skin cell fibroblasts, which have basally increased levels of STAU1, CHOP and p-eIF2*α*, knockdown of *STAU1* by RNAi normalized the levels of UPR mediators (17). We also performed a number of genetic interaction studies in which we crossed SCA2 or TDP-43 transgenic mice with *Stau1^+/-^* mice. ATXN2-Q127 mice haploinsufficient for *Stau1* had improved rotarod performance, improved expression of several cerebellar health-indicator genes including *Pcp2*, and no evidence of Stau1/ATXN2-positive inclusion bodies in Purkinje cells that were observed in ATXN2-Q127 mice (1). ATXN2-Q127 mice either haploinsufficient or null for *Stau1* also had improved or normalized protein abundances for each of mTOR, p-mTOR, p-S6k, p62, LC3 and cleaved CASP3, as did *Thy1*-TDP-43 transgenic mice haploinsufficient for *Stau1* (4). These genetic interaction studies demonstrate that autophagy and neuronal death phenotypes in multiple mouse models can be improved by lowering STAU1 abundance by only 50%.

### Development of a *STAU1* ASO therapeutic

The first step in ASO therapeutic development can take different approaches, with the most extreme being the screening of every possible ASO to a target gene pre-mRNA that could number into the 1000s depending on the pre-mRNA length. We took a conservative but effective approach whereby we determined all possible ASOs to the *STAU1* cDNA then applied criteria for eliminating ASOs with features indicative of an ineffective therapeutic, and of these we selected 118 nearly uniformly distributed ASOs for screening. Our objective was to identify regions in *STAU1* vulnerable to MOE gapmer ASOs supporting RNase H1 activity, revealing lead ASOs that could be optimized by ASO medchem in future research toward developing a *STAU1* ASO for use in patients. Upon performing the primary screen in HEK-293 cells, then retesting efficacious ASOs in SCA2 patient fibroblasts, we arrived at the most efficacious 10 ASOs that we screened for efficacy in BAC-STAU1 mice for lowering STAU1 abundance and restoring the expression of autophagy marker proteins. Along the way, with the production of a new BAC-STAU1 mouse model, we were able to verify abnormal abundance of autophagy proteins (mTOR, p-mTOR, p-S6K, p62, LC3-II) and gene expression (*Calb1, Rgs8, Pcp2, Pcp4, Fam107b*) in cerebellum, further supporting STAU1 role in pathogenesis. Ultimately, we arrived at four of the most efficacious ASOs (ASOs 249, 256, 270, 319) that also had few or no off targets in CNS tissues determined by aligning ASO sequences to the human genome. We defined vulnerable regions in *STAU1* to MOE gapmer ASOs when STAU1 levels are less than 50%. The longest of these is 39 bp in length, defined by ASOs 246-249, including the lead ASO-249. We also identified two ASOs (ASO-045 and ASO-308) that target mouse *Stau1* as well as human *STAU1*, useful for *in vivo* proof of concept studies. Among the top 10 leads ASO-308 had the lowest IC50, however it also had only one mismatch to *SORL1* which has LOF mutations in Alzheimer’s disease and thus will not be considered for human use.

### Target specificity and cytotoxicity

On- and off-target methodologies have limitations. Ultimately, it is typical that the only solution to minimizing off-target RNA-dependent activity is confirmation by empirical preclinical safety testing. In our initial cytotoxicity analyses, we have observed no increases of *Aif1* or *Gfap* in BAC-STAU1 mice treated with any of the top 10 lead ASOs (Fig. 4), except for ASO-256 in a secondary study (Fig. 5) and ASO-308 significantly elevated *Gfap* in wildtype mice in a separate experiment, albeit very slightly in magnitude (Fig. 6), resulting in the elimination of these two ASOs from further consideration. We have also made additional initial efforts to avoid on- and off-targets, described below in this section.

#### Off-targets

To reduce the potential for RNA-dependent off-targets, we evaluated the most efficacious ASOs from our screen for tracts of sequence identity to genes expressed in the CNS. We aligned ASOs to the hg38 human genome. In addition, the *STAU1* ASOs in our study will not target the *STAU2* gene because we verified lack of sequence identity in the targeted regions. Another form of target specificity is when an ASO targets the incorrect splice variant. We have confirmed that all of the 10 most efficacious ASOs (those used in Figs. 3 and 4) would target all *STAU1* isoforms annotated in Ensembl.

#### On-targets

*Stau1^-/-^* mice are viable and have no overt neurological phenotype. The Kiebler group demonstrated *Stau1^-/-^* mice to have no Morris water maze phenotype and mildly reduced open field locomotor activity (7). Also, mice null for *Stau1* showed reduced numbers of functional synapses in hippocampal neurons (7). From the extant literature, it is not clear if this is also observed in *Stau1^+/-^* mice and if this is due to developmental or later functional defects. In our own efforts to evaluate the effect of *Stau1* knockout on open field behavior we did not see reduced locomotion in either *Stau1^-/-^* or *Stau1^-/-^* mice, and we confirmed the observed normal center crossing phenotype (Fig. 9). To evaluate the effect of lowering *Stau1* expression with an ASO therapeutic we also performed open field testing on wildtype mice treated with ASO-45, showing no significant changes in locomotion (Fig. 9).

### *In vivo* proof of concept

To support targeting Staufen1 for treating neurodegenerative diseases we utilized ASO-45 in *in vivo* proof of concept studies. As stated above, we demonstrated that ASO-308 also targeted mouse *Stau1* but ASO-308 also slightly elevated *Gfap* in wildtype mice, but not ASO-45 (Fig. 6). While ASO-308 was highly potent compared to other leads by IC50, in wildtype mice it was equally effective to ASO-45 for lowering mouse *Stau1*. Previously we demonstrated that ATXN2-Q127 mice had reduced intrinsic Purkinje cell firing frequency that was restored by ICV treatment with an ASO targeting *ATXN2* (8). We repeated the experiment in ATXN2-Q127 mice but using ASO-45. We were able to replicate the reduced firing frequency in ATXN2-Q127 mouse PCs in extracellular recordings, and we showed that the firing frequency was significantly increased in ATXN2-Q127 mice treated ICV with ASO-45 (Fig. 7). The firing frequency in this experiment was not fully restored which may be due to ASO-45 not being the most optimized for lowering mouse *Stau1*. In a second *in vivo* proof of concept study, we utilized *Thy1*-TDP-43 mice, in which we previously showed have overabundant Stau1 abundance (3). We treated mice with ASO-45 and demonstrated that the normalized STAU1 abundance, comparable to that found in wildtype mice, resulted in completely normalized ChAT and NeuN abundance in *Thy1*-TDP-43 mouse spinal cord (Fig. 8).

Effective therapeutics are needed for treating neurodegenerative disorders. In the past decade, ASO therapeutics have emerged as viable therapeutics for SMA and DMD and other clinical trials involving ASO therapeutics are underway for neurodegenerative diseases. We have demonstrated that STAU1 is overabundant in cells and animal models relevant to multiple neurodegenerative diseases, including SCA2, ALS, AD, and HD, and that lowering *Stau1* in mouse models relevant to SCA2 and ALS by merely 50% improves disease phenotypes. Future research will be needed to optimize *STAU1* ASO sequence and chemistry toward developing *STAU1* ASOs for treating ALS and potentially other disorders in which STAU1 is overabundant.

## ACKNOWLEDGMENTS

We thank Gentrie Maag and Erika Aoyama for contributing to work in this study. We also extend our gratitude to Siew Peng Ho, William Murray, and Simon MazzaLunn who served as Harrington Discovery Institute (HDI) advisors to DRS in association with support by the HDI Rare Disease Scholarship that he received to develop *STAU1* ASOs. This work was also supported by National Institutes of Health (NIH) / National Institute of Neurological Disorders and Stroke (NINDS) grants R01NS097903, R56NS033123, R37NS033123, R35NS127253, R61NS124965.

---

**Supplementary Data Fig. 1.**
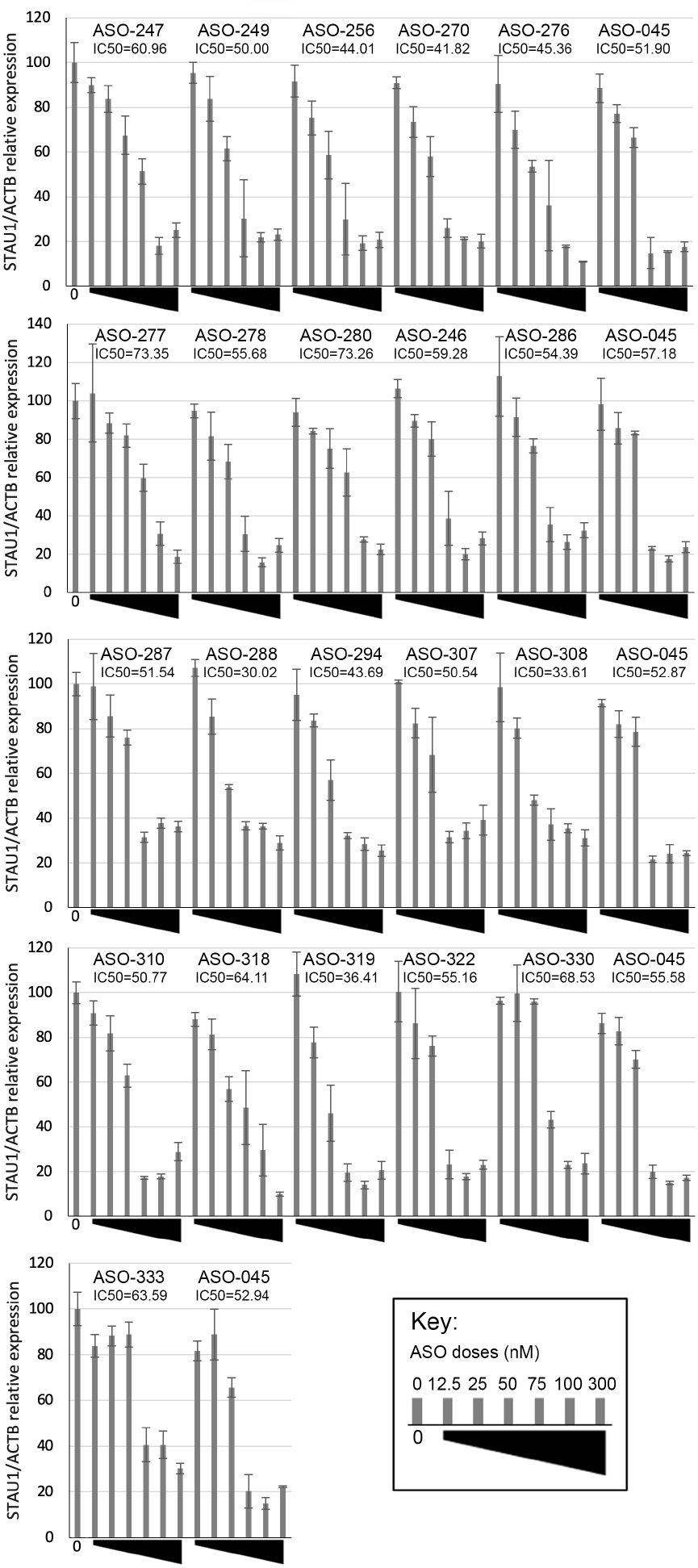
Dose response for lowering *STAU1* expression for 22 ASOs in SCA2 (Q22/Q45) patient derived fibroblast line 500-1. Cells were plated in 384 well plates and triplicate wells were included for each ASO dose. STAU1 abundance relative to *ACTB* was determined by qPCR following 48 hours transfection time. Experiments were done in batches with ASO-045 retested on each plate as a comparative control. IC50s were determined using the Hill equation. The value shown for the 0 nM dose is the mean across the five batches. The IC50 mean and SD for ASO-045 is 54.09 ± 2.2 nM.

